# Do pneumococcal conjugate vaccines (PCVs) reduce childhood pneumonia mortality? An assessment across socioeconomic groups in Brazil

**DOI:** 10.1101/270637

**Authors:** Cynthia Schuck-Paim, Robert J. Taylor, Wladimir J. Alonso, Daniel M. Weinberger, Lone Simonsen

## Abstract

**Background:** Understanding the real-world impact of pneumococcal conjugate vaccines (PCVs) on pneumonia mortality is critical, given the expectation that PCVs can substantially reduce the burden of pneumonia deaths in children under five years. However, surprisingly few post-vaccine introduction studies have estimated the benefit of PCVs for childhood mortality, and results have been inconsistent.

**Methods:** We investigated the long-term trends in child pneumonia mortality in Brazil (1980-present) and assessed the impact of PCV10 on childhood pneumonia mortality, both nationally and in municipalities stratified by socioeconomic status (SES), after the vaccine was introduced in Brazil in 2010.

**Findings:** Between 1980 and 2010, a period when Brazil’s Human Development Index (HDI) rose from 0.55 to 0.71, national pneumonia mortality in children under five decreased 10-fold. Despite rapid uptake of PCV10 following its introduction in 2010, our primary analytical method found no significant decline in national childhood pneumonia mortality, although a secondary analysis found a 10 percent decline in some but not all strata. However, at the municipal level we found significant reductions in childhood pneumonia mortality of up to 24% in low SES strata.

**Interpretation:** Contrary to expectations, we found that PCV use led to at best modest savings in childhood pneumonia mortality at the national level in a middle-income country. In contrast, we found evidence that PCV led to larger reductions in low-income settings; a similar benefit might occur when PCVs are introduced in other low-SES settings. The long-term findings underscore that improvements in nutrition, hygiene, education, and healthcare play a major role in reducing pneumonia mortality.

**Funding:** This work was funded by a grant from the Bill & Melinda Gates Foundation (OPP1114733). DMW also acknowledges support from the Bill and Melinda Gates Foundation (OPP1176267) and the National Institute of Allergy and Infectious Diseases (R01AI123208)

## RESEARCH IN CONTEXT PANEL

### Evidence before this study

Over the past two decades pneumococcal conjugate vaccines (PCVs) have been introduced in more than 100 countries to reduce the burden of pneumococcal disease. Post-marketing effectiveness studies have shown that PCVs reduce hospitalizations for both invasive pneumococcal disease and pneumonia. A reduction in childhood pneumonia mortality, however, is what organizations charged with implementing PCV programs most expect to achieve. For example, WHO has stated that PCVs should be a priority in all countries, especially in those with the highest under-5 pneumonia mortality rates.

Despite such hopes, however, the peer-reviewed literature on PCV-related mortality reduction is surprisingly thin. We searched Pubmed, Google Scholar and Scielo for peer-reviewed post-marketing studies on the impact of pneumococcal conjugate vaccines (PCV) on pneumonia mortality using the terms “pneumonia”, “pneumococcal vaccine/s”, “conjugate vaccine/s”, “PCV”, “pneumococcal”, “impact” and “effectiveness”. We included studies analyzing pneumonia mortality data before and after PCV introduction for any pediatric age group up to December 2017, and excluded studies limited to in-hospital mortality. We identified only five post-marketing effectiveness studies; of these, three were set in a single sub-national population group. Results were inconsistent, with benefits ranging from negligible to extremely large. We therefore set out to assess the impact of PCV10 on childhood pneumonia mortality in Brazil, a large country with high national vaccine coverage and substantial disparity in wealth, allowing comparisons of effectiveness among children from different socioeconomic groups.

### Added value of this study

Our findings show between 1980 and 2010, when PCV10 was introduced, childhood pneumonia mortality had dropped 10-fold in Brazil, concomitant with socio-economic development that led to improved education, sanitation and public health. Our analysis indicates that introduction of PCV10 produced a modest 10% reduction in childhood pneumonia deaths over a three-year period (2011-2014) despite high vaccine uptake. However, after stratifying by socioeconomic status, we found a reduction of up to 24% for the subpopulation living in municipalities characterized by poverty or low maternal education.

### Implications of all the available evidence

Our results indicate that PCV use produced only modest reductions in childhood pneumonia mortality in Brazil, a middle-income country with a near-universal public healthcare delivery system; that result is in jarring contrast to widely held expectations that PCVs will reduce by all-cause childhood pneumonia mortality by about 25% regardless of where they are used. In economically advanced countries—including those that have become so only recently—pediatric pneumonia mortality will likely have already dropped due to general improvements in nutrition, hygiene and medical care, and so the mortality benefit of the vaccine will be smaller. In the world’s poorest countries, however, substantial vaccine-related gains are possible. Confirming that will be difficult, however, because in poorer countries where the pneumonia death toll is still great, good vital statistics data are largely unavailable.

## INTRODUCTION

Despite substantial progress in prevention and treatment, pneumonia is still the leading cause of infectious disease deaths in children under 5 years, causing between 812,000 and 1.1 million deaths in children annually (1). Of all pneumonia deaths in children, 56% are thought to be caused by *Streptococcus pneumoniae* (2), a pathogen that commonly colonizes the nasopharynx of healthy children worldwide. The burden of pneumonia is in addition to the well-characterized burden of otitis media and invasive pneumococcal disease (IPD), which includes cases of septicemia, and meningitis.

Pneumococcal conjugate vaccines (PCVs) have recently been introduced globally to reduce the morbidity and mortality burden of pneumococcal disease. The first formulation, PCV7, was introduced in 2000 and covered seven pneumococcal serotypes. Higher valency versions, PCV10 and PCV13, are now in use, covering 10 and 13 serotypes respectively. Administered to infants in a multi-dose schedule, PCVs are included in the national immunization programs of 135 countries (3). Because PCVs are more expensive than most other vaccines routinely recommended by the WHO Expanded Program Immunization—and may increase total EPI program costs by 60-80% (4)—thoroughly characterizing the total burden prevented is doubly important.

Post-introduction studies of effectiveness have shown that PCVs reduce hospitalizations for both IPD and pneumonia, and provide indirect (herd) protection in highly vaccinated populations to unvaccinated older children and adults (5)(6). The vaccine reduces carriage of covered serotypes throughout the population, and in some countries, vaccine serotypes have been virtually eliminated (7). IPD caused by pneumococcal serotypes not covered by the vaccine generally increases after vaccine introduction; despite setbacks from such replacement. A literature review found a net benefit against IPD in children, whereas in some adult age categories, increases in non-vaccine serotypes completely offset IPD declines in vaccine-targeted serotypes (7).

Reduction in pediatric mortality due to pneumonia, however, is the most widely publicized goal of the global effort to introduce pneumococcal vaccines. For example, to reach Millennium Development Goal 4 (reduce mortality among children under five by two-thirds between 1990 and 2015), WHO says that PCV use could reduce childhood pneumonia mortality by up to 30% (8) and urges that PCVs be a priority in all countries, especially in those with the highest under-5 mortality rates (9). It may stand to reason that fewer children will die from pneumonia if fewer children get pneumonia, surprisingly few published studies have estimated the benefit of PCVs for childhood pneumonia mortality after PCVs have been introduced.

We therefore set out to assess the impact of PCV10 on pneumonia mortality in Brazilian children. Brazil is an ideal settting for such an evaluation. Because PCV10 coverage rose rapidly after its 2010 introduction benefits should become apparent quickly. Brazil has long time series of detailed record-level vital statistics data available. And Brazil has substantial geographic disparity in wealth, allowing comparison of effectiveness among children from different socioeconomic groups; such an analysis should shed light on PCV benefits in poor countries where the pneumonia burden is large but vital statistics data are largely unavailable.

## METHODS

### Mortality data

We accessed publicly available ICD-coded mortality data from the Mortality Information System of the Brazilian Ministry of Health (10). Our age groups for analysis were children aged 3-11, 3-23 and 3-59 months. Infants aged 0-2 months were excluded to avoid perinatal causes of mortality—a major source of childhood mortality in Brazil (11) not preventable by PCV.

The primary outcome for all analyses was all-cause pneumonia (ICD10 J12-18), which in Brazil’s mortality system represented two-thirds (65%) of all deaths with respiratory disease (ICD10 J00-99) as the primary cause of death among children in 2010 (10). We also created time series of all ICD10 subchapters for the period from 2004 (four years after introduction of Hib vaccine in Brazil) to 2014; we used these to control for trends unrelated to PCV use (12). Subchapters with codes that could potentially be affected by PCVs were excluded from the analyses (Supplementary Material). As a comparator outcome, we also analyzed a time series of all deaths that PCV could not be expected to preven (“non-PCV-preventable deaths”), defined as all deaths except those coded as respiratory (any ICD10 J code), otitis media (ICD10 H65-66), sepsis (ICD10 A40-41), meningitis (ICD10 G00-03 and A39), unspecified bacterial infection (ICD10 A49) and encephalitis (ICD10 G04).

Data on births and deaths are systematically collected throughout the year in Brazil, and covered approximately 95% of the population in 2010 (13). Because completeness of vital registration likely increased over time, we conducted sensitivity analyses using mortality time series adjusted for underreporting (Supplementary Material; Figure S1). We found that adjusting the data for under-reporting of deaths made little difference in the point estimates or CIs produced, and did not affect any of the trends observed or conclusions.

### Socioeconomic status (SES) indicators

Socioeconomic conditions are heterogenous across Brazil. Although the southern regions are generally wealthier, poor and wealthy municipalities can be found across the country. Thus, to study the effect of the vaccine in various socioeconomic settings, we stratified Brazil’s 5,570 municipalities into low, medium and high socio-economic categories based on three indicators. We first accessed data on the Human Development Index (HDI) for 2010 (14), a composite measure of income, education and longevity (Figure S2). Because vulnerability to pneumonia and access to treatment can also vary with other SES factors not easily captured by a general index such as HDI, we also stratified the municipalities using two other indicators more closely associated with childhood health: the proportion of children living in extreme poverty (monthly per capita income <$40 US (15), and the proportion of mothers with no primary education, both recognized as important risk factors for severe pneumonia in children (16). For HDI, we formed groups based on UN-defined categories for countries: very low/low (< 0.60), medium (0.60 to < 0. 70), and high/very high (≥ 0.70). For percent children in poverty, we defined low SES as >27 percent of children from the municipality living in extreme poverty, high SES as <3 percent. For maternal education, we defined low SES as >30 percent with no primary education, and high SES as <10 percent. Details of the stratification procedure are available in the Supplementary Material.

### Demographic data

Population estimates for each municipality and age were obtained from the Brazilian Institute of Geography and Statistics (censuses 1991, 2000 and 2010). Monthly population data for each of the SES strata and regions were calculated by cubic polynomial interpolation of the age- and municipality-stratified census data using Popweaver (17), a freely available interpolation software package.

### PCV10 uptake

Coverage data were obtained from the National Immunization Program, that tracks the number of doses administered for each age group in each municipality and year (18). The denominator for vaccine uptake was the number of children in the respective age band, estimated from monthly municipality-specific live-birth data (19). A cohort model that computes coverage in aging cohorts of vaccinated infants from the numbers of doses administered in the post-vaccination period (20) was used to estimate the proportion of children aged 6-23 months who received 3+ doses or the age-appropriate vaccine over time (e.g., one catch-up dose after age one if not eligible for full series).

### Statistical analyses

We have previously described the synthetic control (SC) method in the context of assessing the effect of PCV10 on pneumonia hospitalizations (12). The method uses a Bayesian regression model to combine time series of disease categories that closely correlate with the behavior of the pneumonia time series in the pre-PCV10 period into a weighted composite (synthetic) time series, with categories more closely resembling the pneumonia time series given proportionally higher weights. Weights are then applied to the corresponding disease categories in the post-vaccine period to generate a composite; the composite is taken to be a counterfactual corresponding to what would have happened in the absence of vaccine. Rate ratios are calculated by comparing observed and counterfactual cases in the post-PCV period. A detailed description of the parameters used is given in the Supplementary Material.

We also compared the synthetic control method with a simple regression in which we adjusted for time trends in the pre-vaccine period and seasonality. With this commonly-used time trend model (TT), the counterfactual in the post-vaccine period is generated by assuming that the prevaccine trends continue into the post-vaccine. To adjust for changes in the volume of deaths we used “non-PCV-preventable deaths” as offset (Supplementary Material).

Because the 2009 A/H1N1p pandemic caused a discontinuity in the pneumonia time series for some age groups, with high peaks of mortality observed just before the introduction of PCV10 in Brazil, the pandemic period in Brazil (April 2009 through March 2010) was excluded from all analyses. The pre-vaccine period thus included April 2004 to March 2009 (5 full years, ensuring all seasons were equally represented), and the post-vaccine period April 2010 to March 2014 (4 full years). As it took approximately two years for PCV10 to reach full coverage, rate ratios were calculated for a two-year (evaluation) period, from April 2012 through March 2014.

The R code and data used for these analyses can be downloaded from GitHub at https://github.com/weinbergerlab/PCV-mortality-Brazil.

## RESULTS

### Rates and long-term trends

Pneumonia mortality in Brazilian children decreased by 90% during 1980-2010, dropping from ~150 to ~15 annual deaths per 100,000 (Figure 1). During these pre-PCV decades, the country transitioned from a lower middle-income to a high middle-income economy, and its Human Development Index (HDI) increased from 0.55 to 0.71. Reductions in annual rates of pneumonia mortality were steepest in the 1980s and 1990s, so that most of the decline since 1980 was achieved before the millennium, and thus preceded the introductions of Hib vaccine in 2000 and PCV10 in 2010.

**Figure 1.**
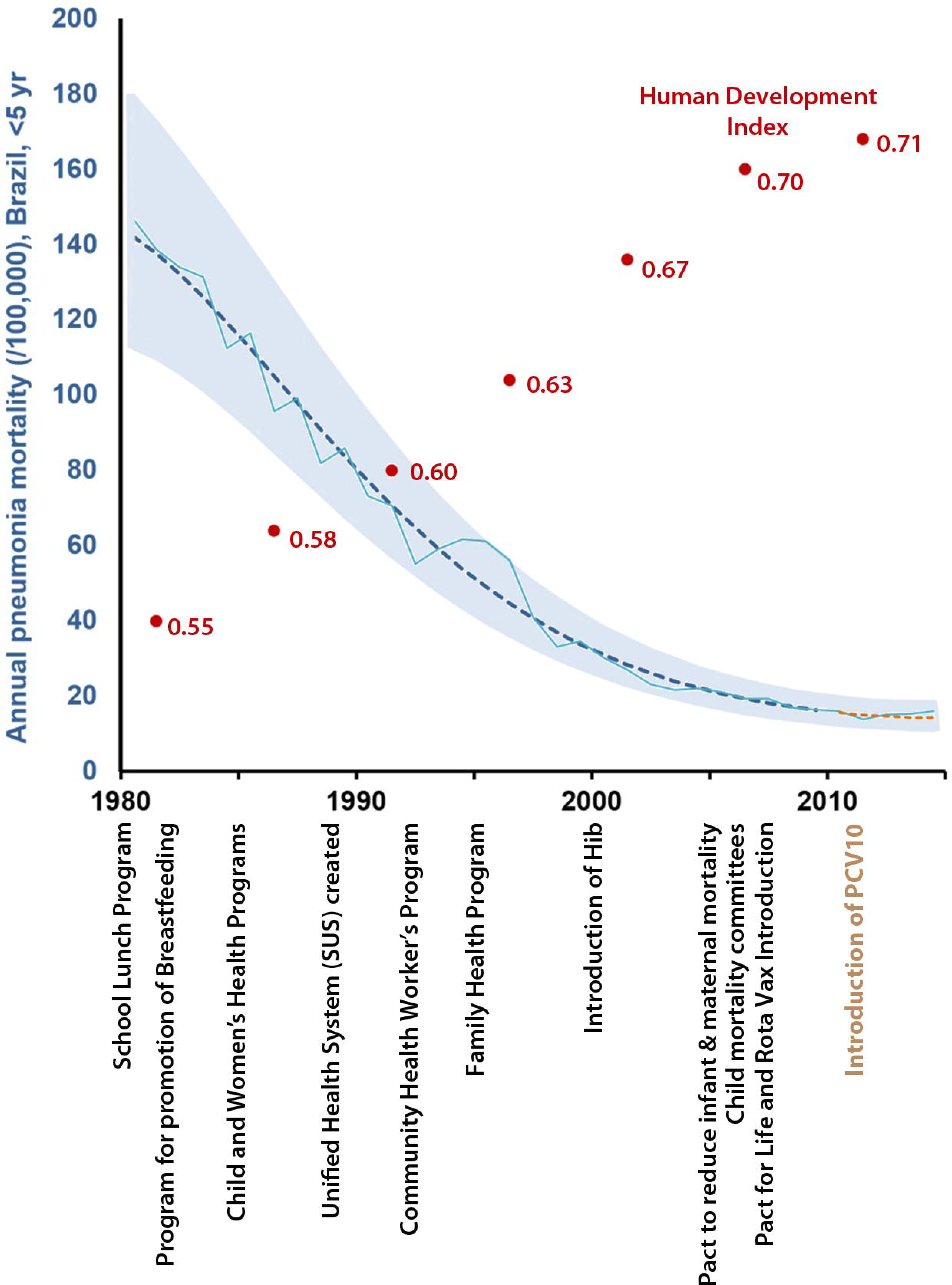
National annual all-cause pneumonia mortality rate in children under five years of age in Brazil, 1980-2014, and Human Development Index (HDI) evolution over time (red dots). Solid line: original pneumonia mortality time series. ICD9-ICD10 change: 1995. In the X-axis some of the major national programs for improving childhood health and reducing mortality are indicated: 1979: National School Food Program (PNAE); 1981: National Program for the Promotion of Breastfeeding (PNIAM); 1984: National Programs for Child Health and for Women’s Health; 1988: Creation of the Unified Health System; 1991: Creation of the community health workers program; 1994: Creation of the Family Health Program (PSF) to increase access to health care in Brazil’s poorest areas; 2000: Introduction of Hib Vaccine; 2004: Pact for Reduction of Maternal and Newborn Mortality; 2005: Creation of local committees for prevention of infant mortality; 2006: Pact for Life (goals included reduction of infant deaths by diarrhea by 50%, and by pneumonia by 20%); 2006: Introduction of Rota vaccine; 2010: Introduction of PCV10.

We also examined raw trends in the years immediately before and after the 2010 introduction of PCV10 (2004 through 2009), using both national data and data from Brazil’s 5750 municipalities stratified by socioeconomic status (SES). During this period, national pneumonia mortality rates declined modestly but were essentially flat thereafter (Figure 2). A similar pattern was observed for non-PCV-preventable deaths (all deaths except those potentially affected by PCV10, Figure S1). We found that the low-SES strata – as based on HDI, proportion of children living under extreme poverty and proportion of mothers without primary school education – had higher rates of pneumonia mortality at the time of PCV introduction in 2010 (Table 1). Pneumonia mortality rates in municipalities stratified by any of the three indeces were relatively flat after 2010 (Figure 2, Figure S3). PCV uptake was lower in the lower SES groups in the first two post-PCV years, but those differences faded by mid-2012 as PCV coverage reached high levels (80-85%) in all strata (Figure 3).

**Figure 2.**
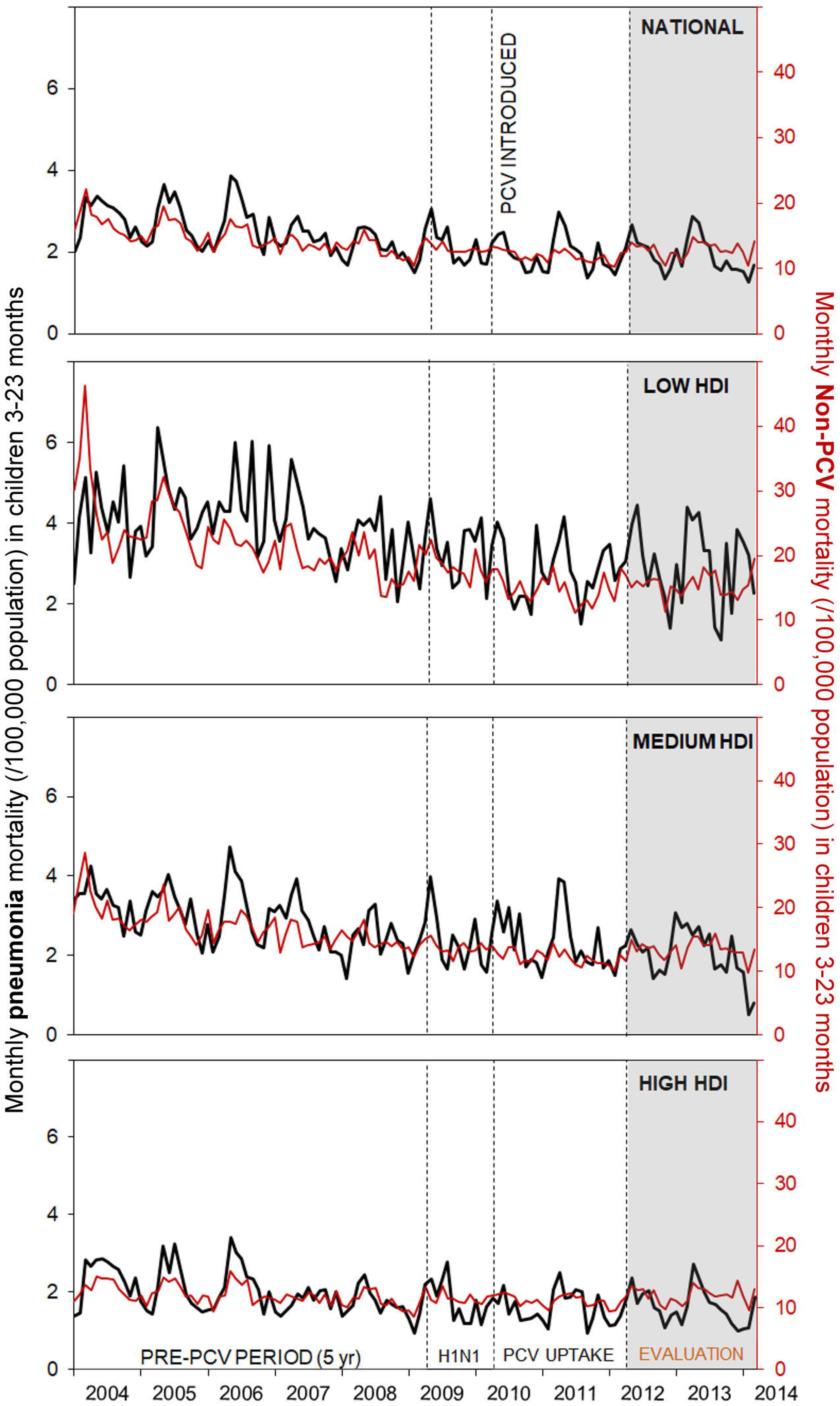
Time series of pneumonia monthly mortality (deaths/100,000 population) among children 3-23 months of age stratified by two indicators of socioeconomic status: percent of children living in extreme poverty (left) and percent of mothers without primary school education (left). The vertical dashed line indicates the time of PCV introduction.

**Figure 3.**
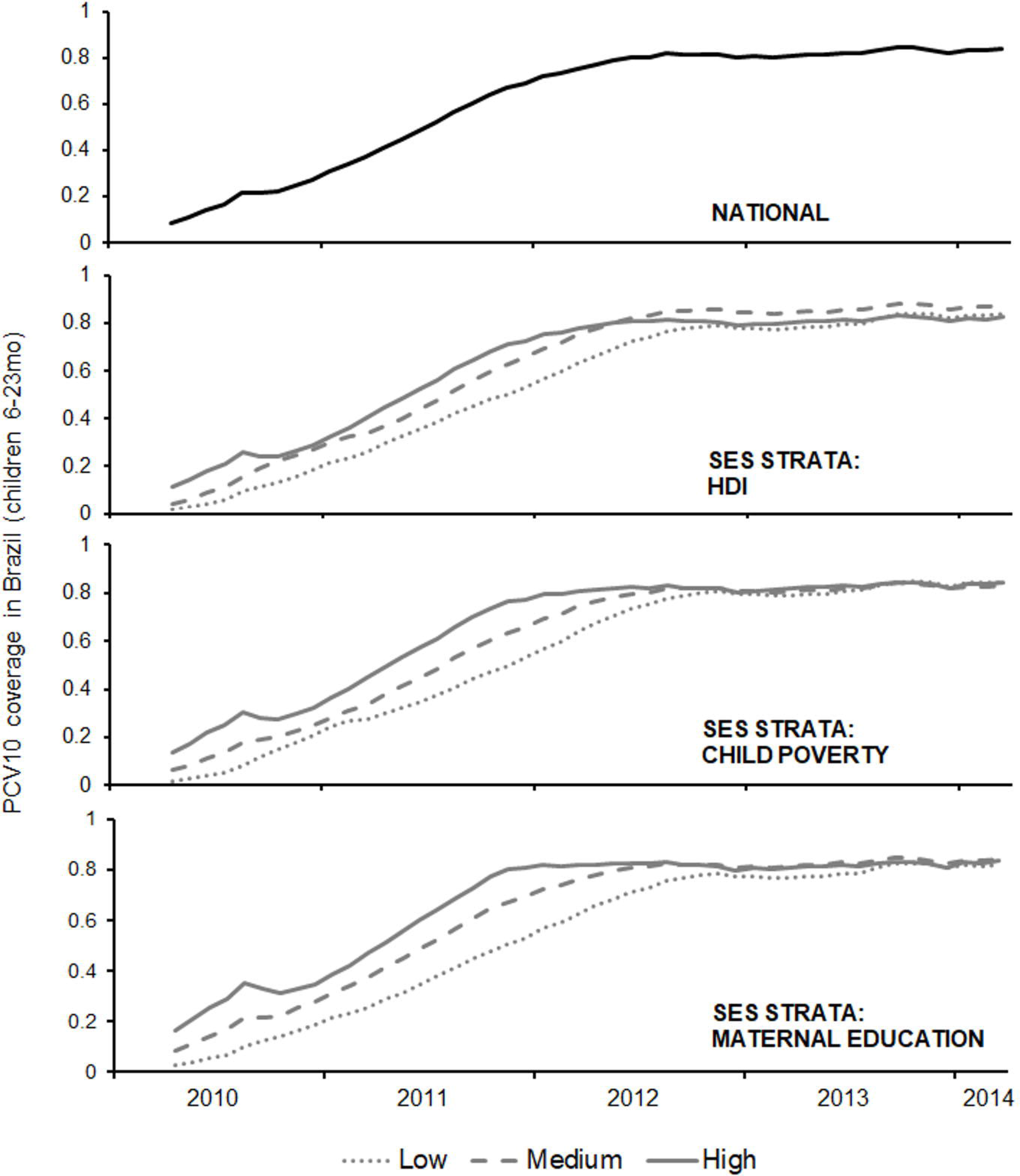
Estimated percentage of Brazilian children fully immunized with PCV10, or who received a catch-up dose of PCV10 for each of the socioeconomic stratifications studies (HDI: human development index; Child Poverty: municipalities with low, medium or high percentages of children living under extreme poverty; Maternal education: municipalities SES scores based on the proportion of mothers without primary school education).

**Table 1.** Annual pneumonia mortality rates per 100,000 (pneumonia deaths /population in 000s) in Brazilian children in 2010, nationally and by municipality sorted by various socioeconomic indicators.

|  |  | 3-11mo | 3-23m | 3-59m |
| --- | --- | --- | --- | --- |
|  | National | 37 (746/2035) | 24 (1126/4730) | 11 (1459/13118) |
| **HDI** | Low | 58 (167/288) | 36 (244/683) | 15 (295/1944) |
|  | Medium | 42 (197/473) | 29 (321/1106) | 13 (413/3097) |
|  | High | 30 (382/1274) | 19 (561/2941) | 9 (751/8077) |
| **Poverty**<br>(based on % children in extreme poverty) | Poorest | 55 (168/305) | 35 (255/720) | 15 (313/2052) |
|  | Medium | 41 (392/961) | 26 (589/2241) | 12 (752/6217) |
|  | Wealthiest | 24 (186/768) | 16 (282/1766) | 8 (394/4843) |
| **Maternal Education**<br>(based on % mothers without education) | Low | 53 (113/213) | 36 (180/504) | 16 (223/1430) |
|  | Medium | 36 (591/1637) | 23 (886/3801) | 11 (1152/10521) |
|  | High | 23 (42/184) | 14 (60/425) | 7 (84/1167) |

### Primary Analysis

We next assessed declines in childhood pneumonia mortality that could be associated with PCV10 use, adjusting for nonvaccine-related trends applying the synthetic control method. At the national level, we observed no significant change in pneumonia mortality in any of the three under-5 age groups, although the point estimates of the relative ratios were all less than one (Figure 4 – black markers; Table S1). When stratified by SES, however, we found decreases in post-PCV10 pneumonia mortality of up to 24% (RR=0.76) in all three pediatric age groups in municipalities with a high percentage of extreme childhood poverty and mothers with no primary education (low SES strata; Figure 4 – black markers; Table S1). A trend toward a higher mortality benefit among successively less privileged groups was most obvious when socioeconomic status was stratified by maternal education, particularly in the two youngest age groups. But such a “dose-response” effect was not apparent when we stratified by poverty or HDI. For HDI, we found a significant decline in mortality only for the 3-23 month age group in the intermediate HDI level. Deaths from non-PCV-preventable disorders, intestinal infectious diseases (A00-A09) and malnutrition (E40-46) were the main variables contributing to the composite of control diseases used to evaluate changes in pneumonia (Table S2); notably, malnutrition (E40-46) was the most heavily weighted variable in most of the lowest SES strata, indicating a correlation between trends in pneumonia and malnutrition in the pre-PC V10 period.

**Figure 4.**
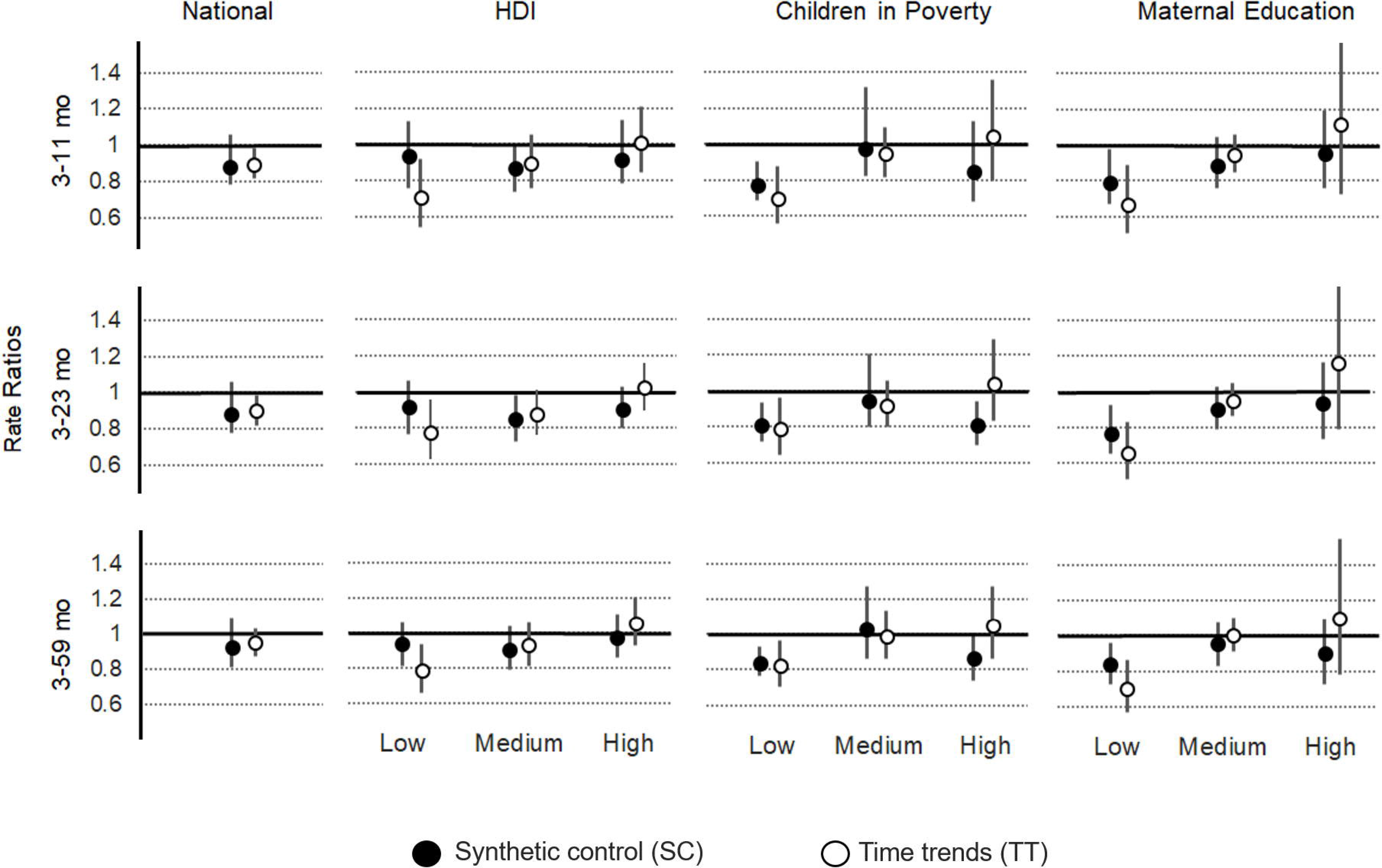
Changes in pneumonia mortality (rate ratios) in three pediatric age groups associated with PCV introduction in Brazil nationally and stratified by three indicators of socioeconomics status (Human Development Index, percentage of children living in extreme poverty, and percentage of mothers without primary school education). Rate ratios were calculated using a model with synthetic controls as the sum of observed deaths from all-cause pneumonia divided by the sum of predicted (counterfactual) deaths during the evaluation period.

### Secondary Analyses

In addition to the primary analyses, we conducted simpler time trends analyses using non-PCV-preventable deaths as the offset to control for previously existing (vaccine-unrelated) trends in the data (see Methods). We observed a pattern similar to the primary SC analysis, but this analysis produced estimated declines in pneumonia mortality in all three “low SES status” strata in all three age groups for which the confidence interval did not overlap “no effect” (Table 2; Figure 4).

**Table 2.**
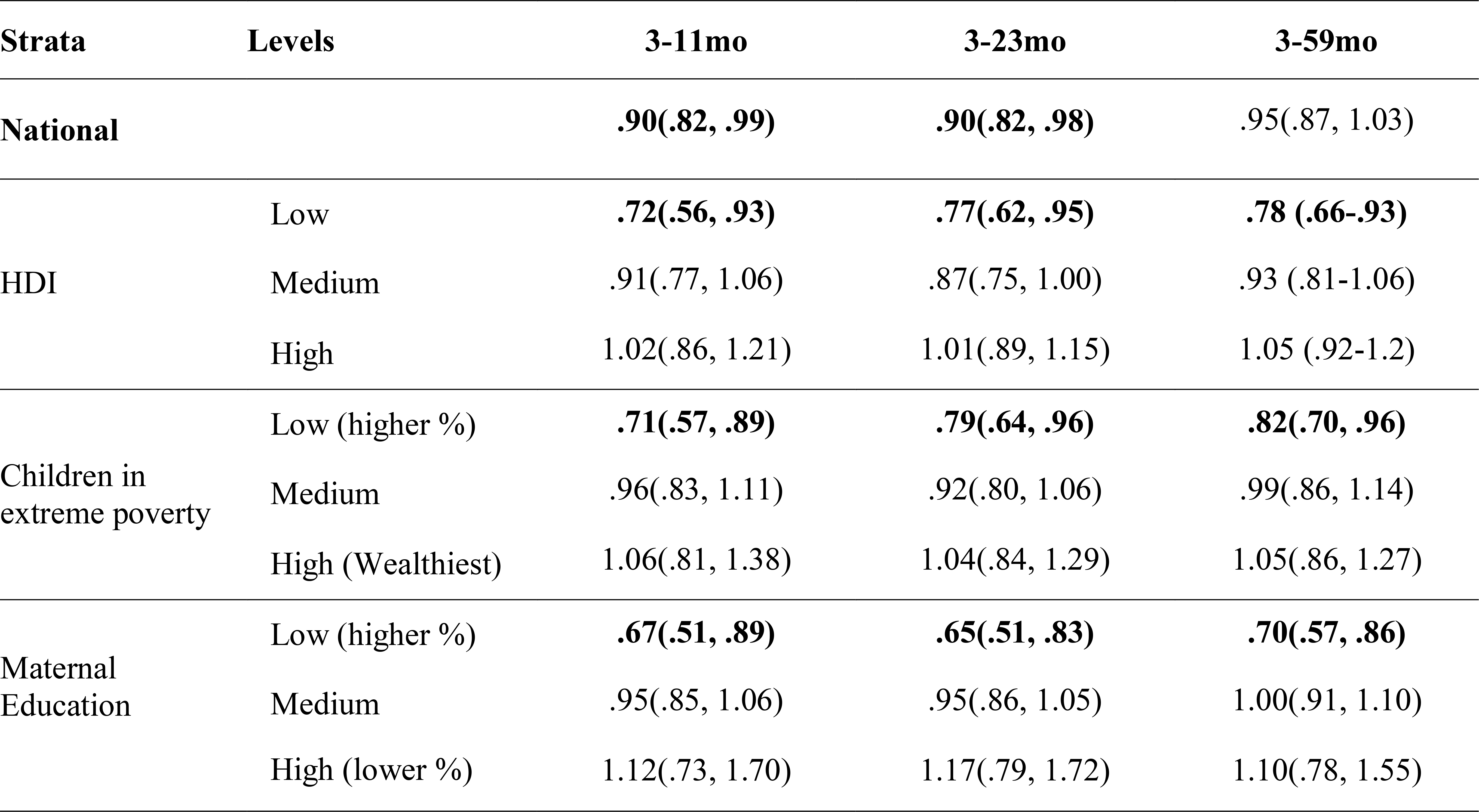
Changes in pneumonia mortality (rate ratios) in three pediatric age groups associated with PCV introduction in Brazil. Analyses were conducted nationally and stratified by three indicators of socioeconomics status, analyzed at the municipality level (Human Development Index, % of children living in extreme poverty, and % of mothers without primary school education). Rate ratios were calculated using a time-trends models adjusted for seasonality and pre-existing trends in the pre-vaccine data and non-PCV deaths as the offset. Rate ratios were calculated as the sum of observed deaths from all-cause pneumonia divided by the sum of predicted (counterfactual) deaths during the evaluation period.

As a validation of the primary analysis, we applied the synthetic control model to a time period when no vaccine effect would be expected, using April 2004 to March 2007 as the training period and April 2007 to March 2009 as the evaluation period. We found that most point estimates were lower than 1, and three low SES strata were associated with significant declines. This potentially indicates still some uncontrolled confounding in these strata. However, credible intervals for the rate ratios for most strata included 1, suggesting no significant change, as would be expected during a pre-PCV period (Figure S4).

## DISCUSSION

We set out to assess the extent to which PCV10 reduced pneumonia mortality in young children after it was introduced in Brazil in 2010. But because Brazil had over many years experienced tremendous economic development, we first looked at trends in childhood pneumonia mortality during 1980-2010, and found that it fell by 90% during these pre-PCV decades (Figure 1). By 2010, pneumonia mortality as a primary cause of death among children aged 3 months to 5 years had leveled off at about 1,500 deaths annually, nationwide, and represented about 12% of total mortality in that age group (Table 1). Such a tremendous improvement represents an unqualified public health triumph. It was surely driven by many factors, including improved housing, nutrition, education and access to health care, especially antibiotic drugs; PCV use, obviously, played no part.

We then analyzed data from the decade surrounding the introduction of PCV10 (2004 through 2014), to assess the extent to which this program reduced childhood mortality. Our primary analyses did not reveal any dramatic reduction at the national level. Instead, they showed a nonsignificant reduction of about 10% in national pneumonia mortality in all pediatric age groups. Our secondary analysis using time trend analysis methods produced similar estimated national reductions, although some of these estimates were—barely—statistically significant.

A 10% reduction in childhood pneumonia mortality four years into the program represents a considerably smaller benefit than expected. The WHO, for example, states that global use of PCVs should reduce the burden of mortality due to pneumonia among children by about 25-30 percent (8). A recent global burden assessment found that about half of all fatal lower respiratory tract infections (LRIs) are caused by pneumococcus, and are thus preventable by vaccine (2). And a crude calculation (multiplying the 85% PCV10 coverage in Brazil by an 80% vaccine efficacy in preventing pneumococcal pneumonia incidence and 50% of all pneumonia being pneumococcal) yields a plausible expected reduction of childhood pneumonia deaths of 34%. Our results, however, are not compatible with PCV having led to such a large reduction in childhood pneumonia deaths on a national level in Brazil.

Several possibilities might explain such a modest national childhood pneumonia mortality benefit of PCV10. Firstly, pneumococcal serotype replacement might have offset the reduction in vaccine-serotype pneumonia achieved by vaccination; this has been observed repeatedly for invasive pneumococcal disease (21); it is even possible that other respiratory pathogens replaced the pneumococcus vaccine serotypes (22). Secondly, changes that occurred in Brazil around the same time PCV10 was introduced might have confounded our analysis, despite our best efforts to minimize confounding using the SC modeling approach (12). For example, in 2010 Brazil banned over-the-counter sale of antibiotics, resulting in a documented and substantial reduction in consumption of penicillin, sulfonamide and macrolide antibiotics in the post-PC V10 period (23). If pneumonia deaths increased as a result—a reasonable proposition—that might offset the vaccine’s benefit. Finally, we note that Brazil had universal publicly funded healthcare system well before the vaccine was introduced. This means that most Brazilian children with severe pneumonia were already getting timely treatment in hospitals so that relatively few died, making the mortality benefit of PCV less apparent.

Although our national estimates of the mortality benefit are smaller than many might have expected, our analyses of low-SES pediatric populations indicate that PCV can prevent up to a quarter of childhood pneumonia deaths in economically disadvantaged municipalities. Across two out of three SES stratifications (HDI, childhood poverty, maternal education), two analytical methods (synthetic control and time trends), and three age groups (3-11 months, 3-23 months, 3-59 months), we observed greater reductions in pneumonia mortality in the low SES strata (Figure 4). We speculate that this pattern could be due to poorer children being more likely to get delayed or inadequate treatment, which increases pneumonia mortality risk. Indeed, we note that the largest and most consistent vaccine effect was observed in the stratum with the least maternal education (16). These results suggest that in poor countries, in which the pediatric pneumonia mortality burden is larger and timely access to care and maternal knowledge about disease more limited, the mortality benefits of PCVs might also be larger. On the other hand, the few significant declines in pneumonia mortality observed in the validation analyses for the SC method were all in the low SES groups, indicating that some confounding may have still been present that partially inflated the estimated vaccine effect in these strata.

Our observational study contributes to a surprisingly small body of post-introduction studies of the effect of PCV use on childhood pneumonia mortality (Table 3). The handful of published studies reported either no overall mortality benefit (24) or mortality benefits larger than what could reasonably be expected based on estimates of pneumonia mortality burden. At the high end, a study set in Chile reported a decline of 35% (9-54%) in mortality from all causes after PCV10 introduction in 3-23 month olds (25)—even though pneumonia deaths contributed only ~8% of all deaths in Chile in 2010 (11). A study set in Peru (26) reported an approximately one-third reduction in pneumonia mortality among infants younger than 12 months after PCV10 introduction, but no decline in other pediatric age groups (12-23m, 2-4y). Studies in Nicaragua reported declines of 33% (20–43%) (27) in all-cause mortality in infants 1-11 months two years after vaccine introduction. A similar study conducted in Nicaragua four years after PCV introduction found an even higher all-cause mortality decline among 1-11 month olds, of 44% (23-59%) (28). However, LRI mortality was estimated to represent less than a third of total deaths in that age group (11). The possibility of negative publication bias—whereby studies showing no benefits are less likely to be published—cannot be ruled out either.

**Table 3.** Published post-marketing studies on the impact of pneumococcal conjugate vaccines (PCV) on pneumonia and all-cause mortality in children. Search terms (Pubmed, Google Scholar, Scielo): “pneumonia”, “pneumococcal vaccine/s”, “conjugate vaccine/s”, “PCV”, “pneumococcal”, “impact” and “effectiveness”, and corresponding terms in Portuguese and Spanish. Studies limited to in-hospital mortality were excludedFigure Captions

| **OBSERVATIONAL POST-LICENSURE STUDIES** |  |  |  |  |  |  |  |  |
| --- | --- | --- | --- | --- | --- | --- | --- | --- |
| **Location** | **Vaccine** | **Age**<br>**<24mo** | **Period** | **Method** | **Adj. for**<br>**pre-vaccine**<br>**trends?** | **Control**<br>**Outcome** | **Mortality**<br>**Outcome** | **%**<br>**Reduction**<br>**\*** |
| **Brazil, Santa**<br>**Catarina (24)** | PCV10 | <12mo | 2006-13 | Pre-Post | Yes | No | J13,J15,J18 | NS<br>(-20 – 34) |
| **Chile (25)** | PCV10 | 2-23mo | 2010, 2011 | Case-Control | NA | No | J13-J18 | 72%<br>(9 – 92) |
| **Nicaragua,**<br>**León (27)** | PCV13 | <12mo | 2008-12 | Pre-Post | No | Diarrhea | Clinical<br>pneumonia | 33%<br>(20 – 43) |
| **Nicaragua,**<br>**León (28)** | PCV13 | <12mo | 2008-15 | Pre-Post | No | Diarrhea | Clinical<br>pneumonia | 44%<br>(23 – 59) |
| **Peru (26)** | PCV10 | <12mo | 2006-12 | Pre-Post | Yes | No | J12-J18 | 35%<br>(9 – 54) |
| **RANDOMIZED CONTROLLED TRIALS (RCTs)** |  |  |  |  |  |  |  |  |
| **Location** | **Vaccine** | **Age** | **Mortality**<br>**outcome** | **Events/Control** | **Events/Vaccine** | **Relative Risk**<br>**(95% CI)** |  |  |
| **South Africa**<br>**(32)** | PCV9 | <24mo | LRI | 160/19,914 | 161/19,992 | NS: 1.01<br>(.76 – 1.2) |  |  |
|  |  |  | All Cause | 242/19,914 | 229/19,992 | NS: 0.95<br>(.79 – 1.13) |  |  |
| **Gambia (31)** | PCV9 | 3-29mo | All Cause | 389/8151 | 330/8,189 | 0.84<br>(.72 – .97) |  |  |
| **L. America (33)** | PCV10 | <24mo | All Cause | 26/11,799 | 19/11,798 | NS: 0.73<br>(.41 – 1.32) |  |  |

\*some authors base estimate on last year in study, others on more than one post-PCV years

Meanwhile, systematic reviews and meta-analysis of randomized controlled studies RCTs (29) (30) did not find that PCVs have a statistically significant effect on pneumonia mortality; this is not surprising given that RCTs are rarely powered to assess deaths and other rare outcomes. One often-cited paper is the PCV9 RCT conducted in The Gambia in the early 2000s, which found that the vaccine reduced all-cause mortality by 16% (3-28%)(31). But this is difficult to reconcile with the fact that only about 12% of all deaths in Gambian children under-5 are due to lower respiratory infectiouns and about 10% from sepsis (11). That implies that a maximum total of 22% of all child deaths were the result of the clinical syndromes caused by pneumococcus, among other pathogens. Assuming that the children enrolled in the trial were representative of the population, PCV would need to prevent more than 70% (16% / 22%) of all LRI, sepsis and meningitis deaths to yield the 16% estimated reduction in all-cause mortality. This leaves very little room for the many other viral and bacterial pathogens that cause respiratory- and invasive bacterial disease mortality.

It is important to know how much PCVs actually reduce childhood pneumonia mortality after they are deployed, and how much the benefit depends on the economic status of the target population. To that end, post-PCV time series analyses similar to ours should be conducted in more countries with high-quality vital statistics data, high PCV coverage and a substantial pneumonia mortality burden. We propose that such studies use the synthetic control method to control for confounding, and exclude infants <3 months of age not yet eligible for PCV. It is important to note that essentially all of the remarkable reduction in childhood pneumonia mortality observed in Brazil after 1980 occurred *before* PCV10 was introduced. That fact underscores the often overlooked power of improved hygiene, health care, education and living conditions that often accompany economic growth, to save lives. Although one cannot know what the impact of PCV might have been in Brazil in 1980, our SES-stratified results suggest the impact would have been considerably greater. These results also suggest that the power of PCVs to save childrens’ lives is greatest in countries that have yet to develop economically, and in which the burden of pneumonia mortality is still high.

## Author Contributions

Conceived of the study: LS, CS, RJT, DW; performed analyses: CS, WJA; Conducted the literature search: CSP; LS, RJT; Wrote the first draft of the paper: LS, CS, RJT; Contributed to and have seen the final draft of the manuscript: CSP, RJT, WJA, DMW, LS

## Acknowledgments

The authors are grateful to the Department of Vital Statistics, a branch of the Brazilian Ministry of Health, for providing the mortality data. We thank Rodrigo Fuentes for information on the dataset from Chile. We thank Stephen Hetterich, Esra Kurum, Joshua Warren and Christian Bruhn for helpful discussions.

